# QRATER: a collaborative and centralized imaging quality control web-based application

**DOI:** 10.1101/2022.12.20.521204

**Authors:** Sofia Fernandez-Lozano, Mahsa Dadar, Cassandra Morrison, Ana Manera, Daniel Andrews, Reza Rajabli, Victoria Madge, Etienne St-Onge, Neda Shafiee, Alexandra Livadas, Vladimir Fonov, D. Louis Collins, Alzheimer’s Disease Neuroimaging Initiative

**Affiliations:** McConnell Brain Imaging Centre, Montreal Neurological Institute, McGill University, Montreal, Quebec, Canada; Department of Neurology and Neurosurgery, McGill University, Montreal, Quebec, Canada; Department of Psychiatry, McGill University, Montreal, Quebec, Canada; Douglas Mental Health University Institute, Montreal, Quebec, Canada

**Author notes:** Data used in preparation of this article were obtained from the Alzheimer’s Disease Neuroimaging Initiative (ADNI) database (adni.loni.usc.edu). As such, the investigators within the ADNI contributed to the design and implementation of ADNI and/or provided data but did not participate in analysis or writing of this report. A complete listing of ADNI investigators can be found at: http://adni.loni.usc.edu/wp-content/uploads/how_to_apply/ADNI_Acknowledgement_List.pdf.

## Abstract

Quality control (QC) is an important part of all scientific analysis, including neuroscience. With manual curation considered the gold standard, there remains a lack of available tools that make manual neuroimaging QC accessible, fast, and easy. In this article we present Qrater, a containerized web-based python application that enables viewing and rating of previously generated QC images. A group of raters with varying amounts of experience in QC evaluated Qrater in three different tasks: QC of MRI raw acquisition (10,196 images), QC of non-linear registration to a standard template (10,196 images) and QC of skull segmentation (6,968 images). We measured the proportion of failed images, timing and intra- and inter-rater agreement. Raters spent vastly different amounts of time on each image depending on their experience and the task at hand. QC of MRI raw acquisition was the slowest. While an expert rater needed approximately one minute, trained raters spent 2-6 minutes evaluating an image. The fastest was the curation of a skull segmentation image, where expert raters spent on average 3 seconds per image before assigning a rating. Rating agreement also varied depending on the experience of the raters and the task at hand: trained raters’ inter-rater agreement with the expert’s gold standard ranged from fair to substantial in raw acquisition (Cohen’s chance corrected kappa agreement scores up to 0.72) and from fair to excellent in linear registration (kappa scores up to 0.82), while the experts’ inter-rater agreement of the skull segmentation task was excellent (kappa = 0.83). These results demonstrate that Qrater is a useful asset for QC tasks that rely on manual curation of images.

## Introduction

The first step in any kind of neuroscientific analysis is data curation. Doing quality control (QC), that is assuring that the data was acquired correctly, and that its quality meets established standards that would permit it to be reliably analyzed, is not a trivial task. Specifically in neuroimaging, it has been demonstrated that motion artifacts introduced in MRI acquisition can compromise the results in structural (Alexander-Bloch et al., 2016; Ducharme et al., 2016; Gilmore et al., 2021), diffusion (Yendiki et al., 2014), and functional (Power et al., 2012) analyses. While the effects of some acquisition artefacts can be minimized in the preprocessing pipeline, other artefacts might not be recoverable and would need to be excluded from the analysis.

QC does not apply only to acquisition; quality must be verified after the different processing pipeline steps. As the amount of data has increased in recent years, QC has become more difficult and time-consuming. For this reason, several automatic QC tools have been developed for MRI data acquisition. Pizarro, et al. (2016) obtained 80% accuracy using a support vector machine algorithm to automatically categorize the quality of 3D-MRI when compared to manual QC. MRIQC (Esteban et al., 2017) is another quality assessment classifier that can automatically extract a vast number of quality metrics from a T1 or T2 structural image and, from this data, yield a binary classification of the image’s quality. In addition, processing steps like tissue segmentation and image registration to stereotaxic spaces require quality control and automatic QC models have been proposed. Qoala-T is a supervised-learning tool that works with Freesurfer’s segmentation and quality metrics output to generate a single measure of scan quality ranging from 0 to 100 (Klapwijk et al., 2019). For image registration, two deep learning approaches have been proposed: DARQ (V. S. Fonov et al., 2022) and RegQCNET (Senneville et al., 2020) with impressive performance.

These automatic tools are still limited in the form of intense computer requirements for model training or processing times. While they are a vast improvement in relation to the time-burden of manual QC and can be used to automatically identify images that are clear FAILures or clear PASSes, they have not yet replaced the need for manual QC for images where the quality is less certain. Indeed, most of these papers suggest some form of human visual inspection to rate the *uncertain* results justifying the need for an efficient tool to complete manual QC.

Manual curation is still the most reliable approach to quality control (Monereo-Sánchez et al., 2021). The conventional method often involves opening an MRI volume in an image viewer and logging the QC result in a separate file. This can be cumbersome, slow, error-prone and makes collaboration difficult.

We developed Qrater (Quality Rater), a web-based python application, to address these limitations. Qrater was designed to enable multiple users to simultaneously perform visual inspection of large databases from both local and remote computers quickly and easily. The closest similar QC tool is the QC System of the Laboratory of Neuro Imaging (LONI), a web-based platform where any user can upload imaging data to aid QC by the automatic generation of quality metrics and visualization of the data for manual curation (Kim et al., 2019). The main difference between our Qrater and the tool from LONI is that the latter was made to be primarily run on the LONI servers where the users have to create an account and upload their data. While they offer the option to run an offline version of LONI locally, it expects the user to have pre-installed a suite of imaging software (FSL, AFNI, Freesurfer and SPM). Qrater, on the other hand, is run from a Docker container locally in the user’s own server or PC and works with previously generated QC images in .jpg or .png formats instead of the DICOM or NIfTI files themselves.

After looking at intra- and inter-observer QC variability using Qrater, we ran benchmark tests on a series of different datasets to test the usability of Qrater within the context of three distinct steps of neuroimaging processing pipelines: QC of raw acquisition, QC of linear registration, and QC of segmentation. We also evaluated the time it would take a team to rate the raw imaging quality of close to 9000 images from ADNI 3 (Weiner et al., 2017). Qrater demonstrated an improvement over our previous manual quality rating methods in relation to both speed and usability.

## Methods

### The application

We set out to develop Qrater, an open-source web-based application for imaging quality control by visual inspection, that would fulfil our specific QC needs. The main design criteria include 1) sufficient, 2) accessible, 3) adaptable, 4) centralized, and 5) collaboration centric. These criteria are detailed below.

#### Sufficient

Qrater is built as a set of software tools in the form of Docker containers. In a single tool, Qrater encompasses both a Graphical User Interface (GUI) where the user can view, search, and inspect image files, and a built-in database where all the quality information (QC status and comments) is stored and managed.

#### Accessible

The use of Docker images makes Qrater easily installable and manageable on any operating system. Built with the purpose of making the QC task as approachable and easy as possible, Qrater can be used by users with minimal or no programming skills. Once installed and set-up, most of Qrater’s features can be accessed through the GUI (see Fig. 1). Viewing and rating the images; loading new datasets; browsing, searching, or exporting the QC data; all of this can be done from the web browser. The GUI also includes several imaging features that facilitate and speed up the visual inspection process: a magnifying glass; built-in image controls for brightness, saturation, and contrast; and key-bindings for navigation and rating.

**Figure 1.**
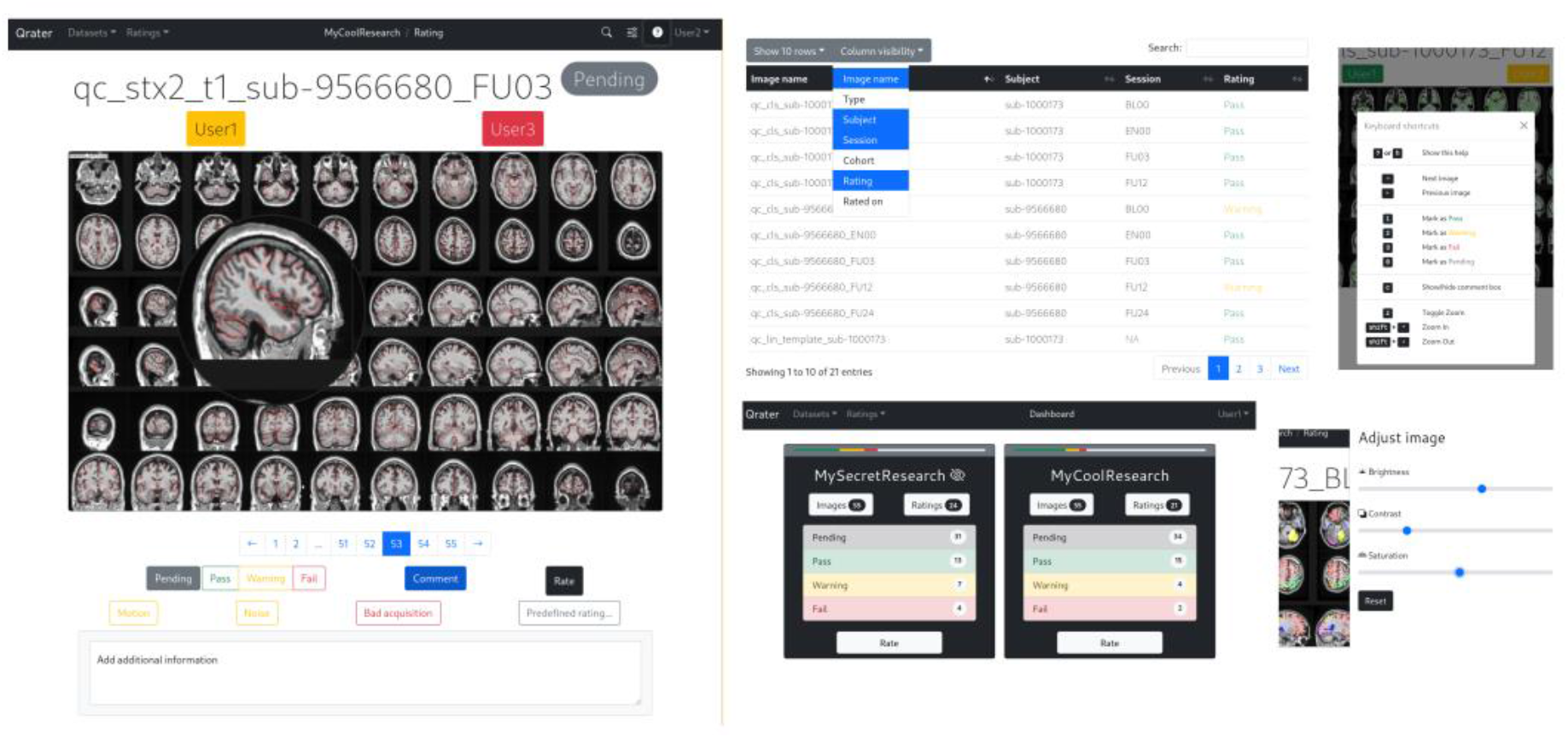
Examples of the main features of Qrater’s Graphical User Interface. Rating window with team’s ratings above the image viewer, magnifying glass, comment box and predefined ratings; database of images and ratings with the ability to toggle columns’ visibility and search; private and open datasets; key-bindings for faster ratings; built-in image editing.

#### Adaptable

While there are other automatic tools designed to work with specific MRI contrasts or to address particular QC tasks like acquisition or image registration, we opted instead to create a general tool that can be easily adaptable to distinct types of QC. Any QC task that can be achieved by the visual inspection of an image in .jpg or .png format can be facilitated by Qrater.

#### Centralized

Whether installed locally on a personal computer or in a High-Performing Computer (HPC) cluster, Qrater can be accessed remotely via SSH by multiple users simultaneously. When installed on a server, data already located in the server can be loaded without the need to copy or duplicate it. This is particularly useful when dealing with large datasets.

#### Collaboration centric

Qrater was built with collaboration in mind. Its database is capable of handling simultaneous requests. Multiple users can visually inspect the same images without interfering with each other while also offering the feature of displaying the QC status given by other raters. All the QC status information can be browsed and searched in the built-in database accessible through the GUI.

### Benchmark Tests

#### Raters

The different datasets were rated by 12 different members of our team with varying ranges of QC expertise; three were expert raters with extensive experience in image processing and related QC, while the other nine were trained following the protocol described below.

#### QC training and intra-, inter-rater consistency

The team of expert raters designed a protocol and developed training material for the QC of MRI raw acquisition and linear registration. The protocol consisted of a detailed description of the criteria for passing or failing an image depending on coverage, noise, motion, and intensity uniformity for raw acquisition; and on symmetry, rotation and scaling for the linear registration (see Table 1). A set of videos was created with instructions and examples of what artifacts to look for.

**Table 1.**
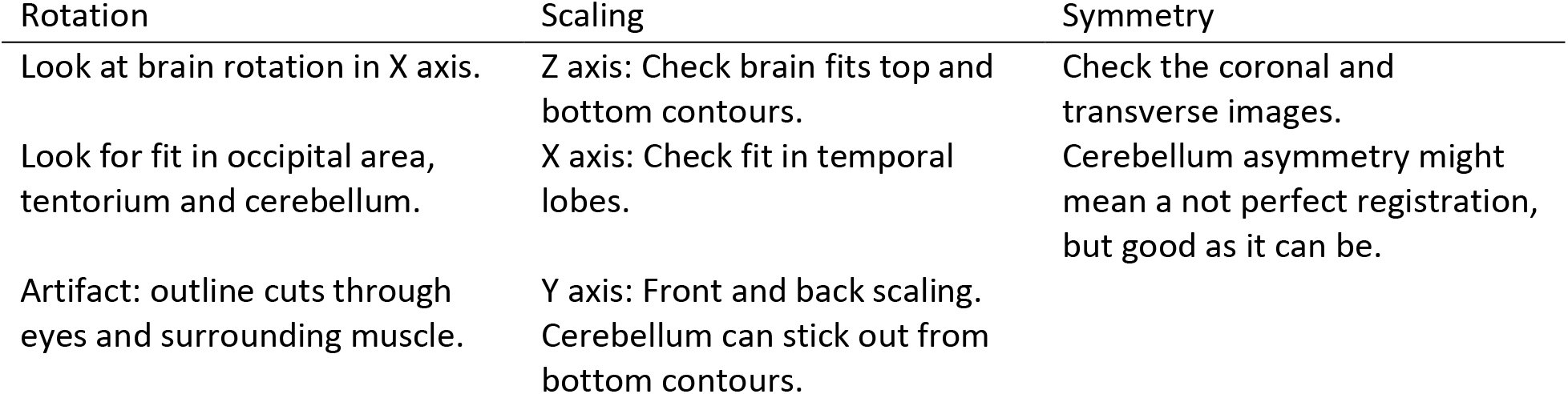
Examples of the instructions given in the Linear Registration experiment.

| Rotation | Scaling | Symmetry |
| --- | --- | --- |
| Look at brain rotation in X axis. | Z axis: Check brain fits top and bottom contours. | Check the coronal and transverse images. |
| Look for fit in occipital area, tentorium and cerebellum. | X axis: Check fit in temporal lobes. | Cerebellum asymmetry might mean a not perfect registration, but good as it can be. |
| Artifact: outline cuts through eyes and surrounding muscle. | Y axis: Front and back scaling. Cerebellum can stick out from bottom contours. |  |

The remaining team members attended two training sessions where the expert raters gave the instructions for the QC task, going through several examples of images that would fail the protocol. This served the purpose of setting a level of strictness that would be similar for the whole group. These sessions were recorded and available to be reviewed later at any time.

To evaluate raw acquisition, for every 3D T1W MRI volume, we created a mosaic image in jpg format containing 60 slices of the brain (20 from the coronal plane, 20 from the axial plane, and 20 from the sagittal plane; see Fig. 2A). To check the quality of the registrations, similarly to the raw acquisition QC task, we created mosaic images of axial, coronal and sagittal slices (see Fig. 2B). Particularly, these images consisted of the subject’s T1w brain MRI, overlaid with the standard brain template’s brain contour. In contrast with the raw acquisition mosaic images, these slices were selected to be at predetermined locations in the stereotaxic brain-based coordinate system to aid the visual inspection.

**Figure 2.**
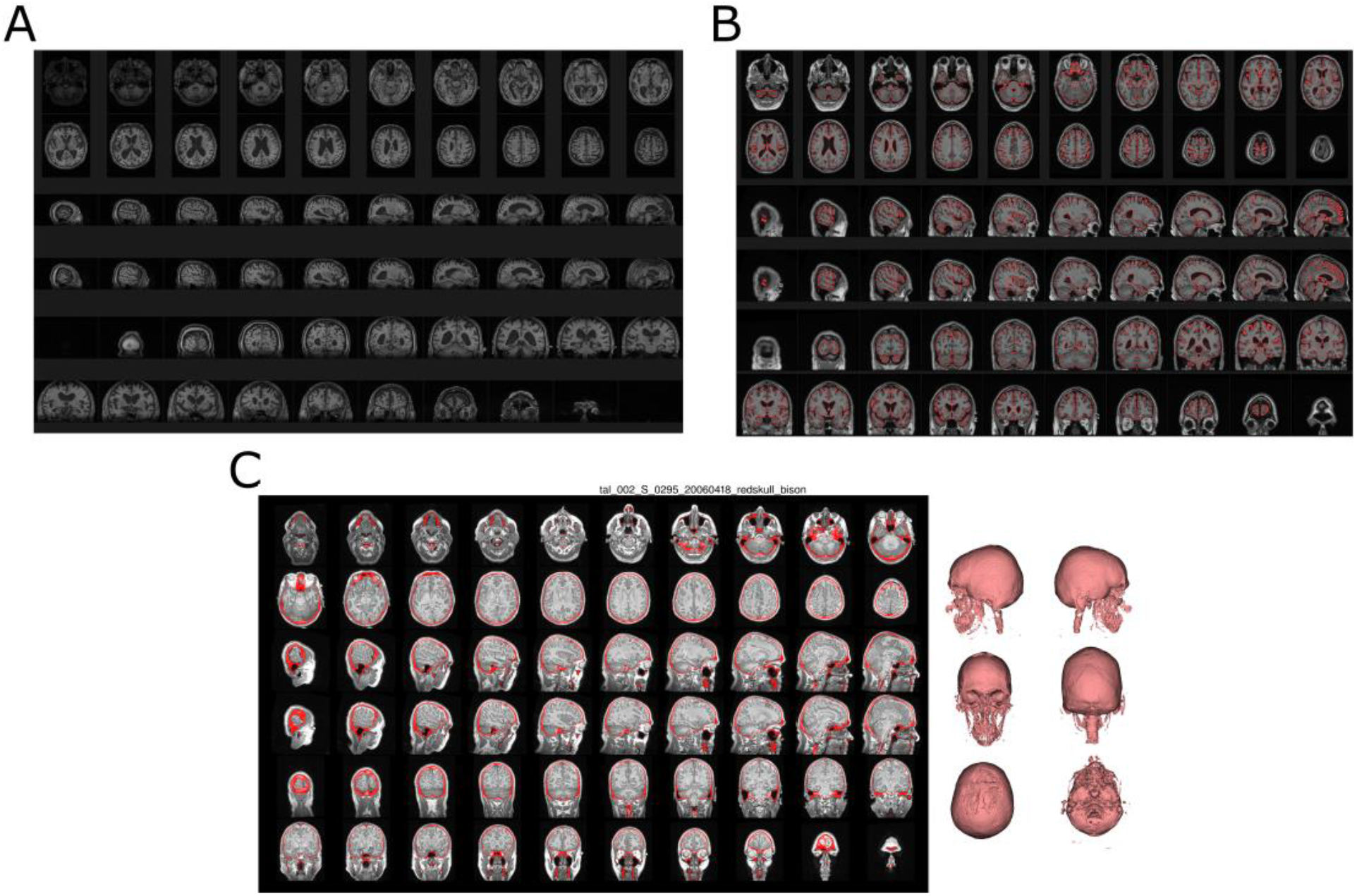
Example images generated for QC: A) mosaic image containing 60 slices of the brain, 20 axial, 20 sagittal, and 20 coronal images; B) 60 slices (axial, sagittal and coronal plane; 20 each) from the subject’s MRI, resampled into the standard Stereotaxic coordinate system, with the standard template’s cortical boundaries overlaid in red; C) an image with 60 slices (axial, sagittal, coronal; 20 each) of the subject’s head with the segmented skull tissue highlighted in red on the left side and six 3D reconstructions of the skull on the right side.

The team of experts (Raters 1 & 12) then pre-selected 99 images previously inspected by them for each of the two QC tasks. These images were particularly chosen to comprise a balanced set of passing and failing images; particularly, 33 clear passes (agreed by both experts), 33 clear fails (agree by both experts), and another 33 borderline cases where the two experts disagreed on the QC status.

The datasets were loaded into Qrater with limited permissions to avoid inter-rater influence.

One of the experienced raters (Rater1) reviewed again the selected images on two separate occasions to estimate the expert’s intra-rater agreement. To minimize learning effects, the two QC sessions were separated by a 2-week delay. Finally, to arrive at a consensus gold standard, Rater 1 re-examined the images that had a different QC status between these two sessions. This consensus process was done for both the raw acquisition and the linear registration datasets. Nine trained raters (Raters 2-10) went through the first set of 99 images to evaluate the training of raw acquisition, while eight raters (Rater 10 dropped out of the exercise) followed up with the rating of the 99 images evaluating linear registration. From the data recorded in these training exercises, we extracted the amount of time required for each rater to QC each image, and we used the ratings to estimate inter-rater reliability with each other and with the expert’s gold standard

Both intra- and inter-rater agreement were assessed with Cohen’s chance corrected kappa coefficient (Landis & Koch, 1977).. It is important to note that by accounting for agreement by chance, Cohen’s kappa is stricter, yielding lower numbers, compared to the Sørensen-Dice coefficient often used to measure agreement in segmentations (e.g., for a PASS/FAIL QC criteria of a dataset of 100 images, if 2 raters agree on 75 of the images, Cohen’s Kappa would be equal to 0.5).

Timing for both the expert’s and trainees’ QC was estimated by looking at successive timestamps for every recorded rating while ignoring time lapses longer than 15 minutes, assuming this was time spent away from the QC task at hand. Given the non-normal distribution of the timing data, we used the nonparametric Wilcoxon rank sum test to compare the timings of the different ratings. Timestamps have a discretization of one second. Once the rating of the datasets was completed by all the raters, the team met again for a follow-up discussion of the disagreement in ratings and to answer any questions that had arisen from the training exercises.

#### QC tasks

Three QC tasks were completed to evaluate how the tool can be used in real-world collaborative projects. The tasks were chosen to be from three distinct stages of a neuroimaging analysis pipeline: data acquisition, pre-processing and analysis derivatives.

##### Raw acquisition QC

The first QC task rated the raw acquisitions of 10,196 3D T1w brain MRI from ADNI3 (Weiner et al., 2017). ADNI data used in the preparation of this article were obtained from the Alzheimer’s Disease Neuroimaging Initiative (ADNI) database (adni.loni.usc.edu). The ADNI was launched in 2003 as a public-private partnership, led by Principal Investigator Michael W. Weiner, MD. The primary goal of ADNI has been to test whether serial magnetic resonance imaging (MRI), positron emission tomography (PET), other biological markers, and clinical and neuropsychological assessment can be combined to measure the progression of mild cognitive impairment (MCI) and early Alzheimer’s disease (AD). Since ADNI data has already gone through QC before being made public, we expected a very high rate of PASSing scans but still looked for issues related to coverage, wrap-around, noise and movement artefacts as our thresholds for these characteristics were different from those of ADNI.

We divided the 10,196 generated mosaic images into eight separate datasets to be reviewed and inspected. A team consisting of the expert rater and seven out of the nine people who had undergone the training and the inter-rater reliability exercise for raw acquisition (raters 9 & 10 did not continue) went through these datasets following the same protocol as before to rate each image as PASS, WARNING, or FAIL. To measure the usability of the application, we timed how long it took to rate each image following the same procedure as the training exercise. We used the non-parametric Kruskal-Wallis test by ranks to see if there was a difference in timing across QC labels and compared the difference in time of the this and the previous training task using the Wilcoxon rank sum test.

##### Registration QC

The second QC task was to review linear stereotaxic registration (Dadar et al., 2018) of 10,196 3D T1w brain MRI from ADNI3 (Weiner et al., 2017) into the space of the MNI-ICBM152-2009c standardized brain template (V. Fonov et al., 2011). All eight raters who completed the training and inter-agreement evaluation (raters 2-9) carried on with this portion of the exercise. For this exercise, we instructed raters to limit their ratings to Pass or Fail.

Just like with raw acquisition, we measured timing following the same procedure described before. Given that this exercise was limited to PASS/FAIL, we used Wilcoxon rank sum test for both evaluation of time difference across the two ratings and between the training and this exercise.

##### Segmentation QC

The third QC task assessed the performance of an image segmentation pipeline that extracted the skull from T1w MRI. Since the main goal of skull segmentation was to identify the external surface of the skull to use as a feature in longitudinal registration, segmentation did not need to be perfect. Success was defined in having at least 95% of the top half of the skull properly segmented (i.e., 2-3 holes of less than 2 cm^2^ were allowed). To make visual evaluation more efficient, 3D renderings of the segmented skull were included with the 2D mosaic of MRI slices overlaid with the segmented voxels in red (see Fig. 2C). Two members of our team with extensive experience in QC used Qrater to visually inspect these QC images and rate them as PASS or FAIL. One (rater 12) went over the whole dataset (N=6,968) and the second one (rater 1) reviewed a subset of these same images (N=1,746) to evaluate inter-rater agreement. Reliability was measured using Cohen’s Kappa. The time spent by the raters on each image was measured the same way as in the other QC tasks and compared using Wilcoxon rank sum test.

## Results

### QC training protocol and agreement

Rater 1, after visually inspecting the same 99 images evaluating raw acquisition at two different timepoints, agreed on 91 of the 99 images (Fig. 3A), thus presenting excellent intra-rater agreement (Landis & Koch, 1977), with a Cohen’s kappa value of 0.81 (Z=8.14; p <0.001). Rater1 re-examined the eight images, and finalized four as PASS and four as FAIL for a final consensus. The resulting gold standard for raw acquisition consisted of 30 FAILs and 69 PASSes.

**Figure 3.**
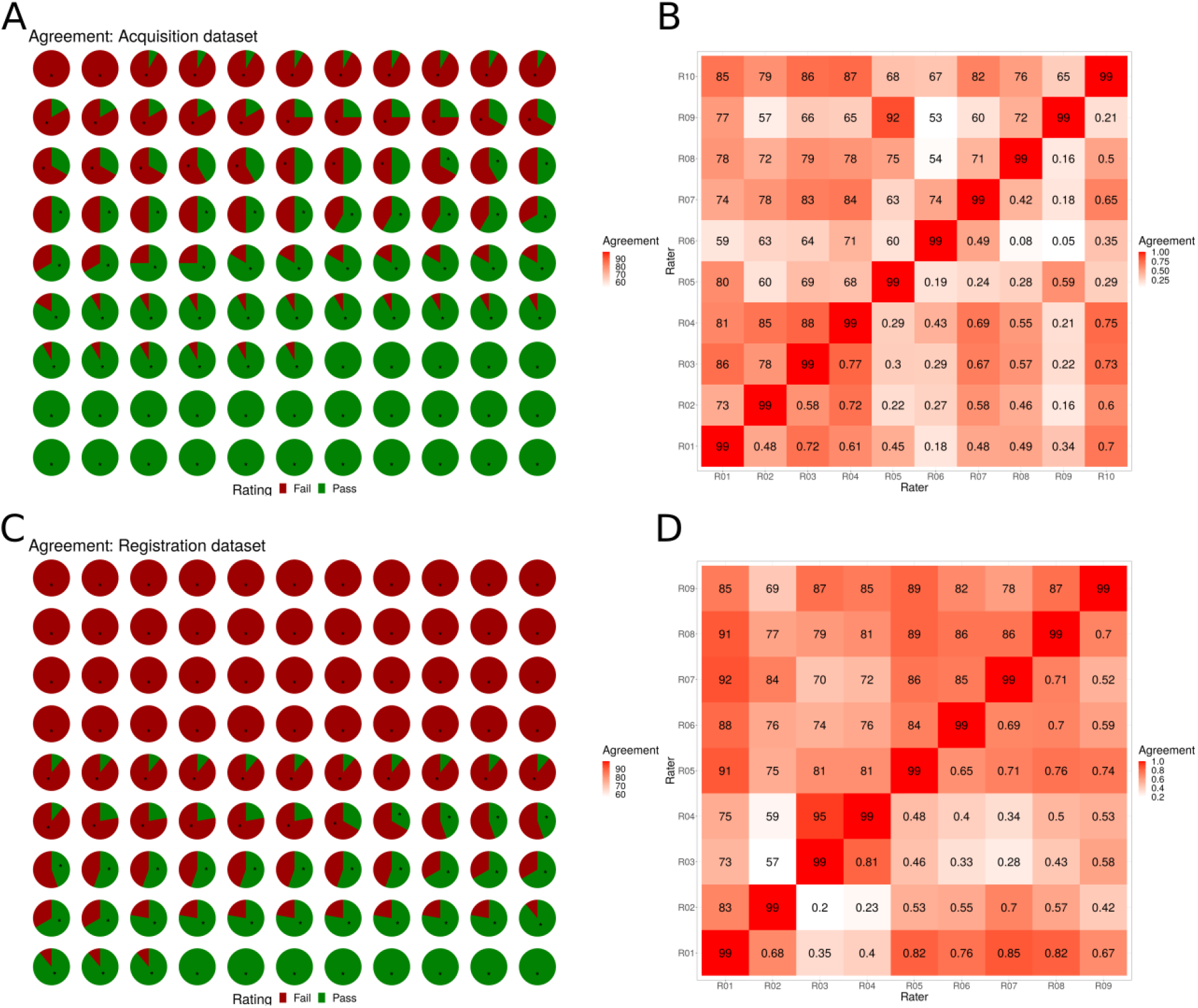
Intra- and inter-rater agreement data from the training exercises. A+C) Pie charts showing relative agreement of the expert consensus rating and the trainees’ ratings, where Red=FAIL, Green=PASS. The first three rows represent training images pre-selected by two experts to have FAIL status. The middle three rows represent images pre-selected where the two experts disagreed. The last three rows represent images pre-selected by two experts to have PASS status. Note however that interrater agreement is based on the consensus PASS:FAIL labeling. Asterisks represent the ratings given by the expert (Rater 01) during the intra-rater reliability evaluation. Panel A shows the results for the raw acquisition QC task and panel C, the stereotaxic registration task. B) Confusion matrix showing interrater agreement for the raw acquisition QC task between the expert’s gold standard (R01) and the trainees (R02-R10) where the upper triangle shows the number of images in agreement and the bottom triangle notes Cohen’s Kappa. D) Confusion matrix showing inter-rater agreement for the linear registration QC task between the expert’s gold standard (R01) and the trainees (R02-R09). The upper triangle represents the number of images in agreement and the bottom triangle notes Cohen’s Kappa.

Rater 1 showed an excellent intra-rater agreement on the linear registration dataset as well, agreeing on 93 of the 99 images (Fig. 3C) with a Cohen’s Kappa value of 0.87 (Z = 8.69, p < 0.001). The reexamination of the 6 conflicting images resulted in 5 FAILs and 1 PASS. The gold standard for linear registration ended up being 62 FAILs and 37 PASSes. For both raw and registration QC tasks, the unbalanced PASS:FAIL proportion is accounted for by using Cohen’s chance corrected Kappa.

In the raw acquisition dataset, 29 of the 99 cases had full agreement amongst all 10 raters (expert’s consensus rating and 9 raters, see Fig. 3A). Of these, all raters agreed on 27 of the 69 cases with a consensus rating of PASS, but only 2 cases with a consensus rating of FAIL. At least eight of the raters agreed on 67 of the 99 cases. Interestingly, none of the preselected *intermediate* cases achieved complete agreement.

In the registration task, raters fully agreed with the consensus labelling in 52 of the 99 cases (44 FAIL and 8 PASS). In 15 cases, only one rater disagreed with the consensus QC status (11 FAILS and 4 PASS) (see Fig. 3C). In the 33 preselected FAIL cases, there was complete agreement amongst raters. Interestingly, raters unanimously FAILed 11 of the preselected *intermediate* cases. However, of the 33 preselected PASS cases, only eight obtained complete agreement.

The detailed inter-rater agreement for both tasks is presented as a confusion matrix with the number of images and the Cohen’s kappa values between each rater in Fig. 3 subsections B & D for the raw acquisition and linear registration tasks, respectively.

The timestamps for the ratings showed that the expert rater time varied considerably between sessions and tasks. On the raw acquisition, Rater 1 took a median of 44 seconds the first session in contrast to 25 seconds the second session (W = 6,770, p < 0.001). By contrast, there was no decrease in timing between the two sessions of the linear registration task (16 seconds vs 15.5 seconds, W = 4,628.5, p = 0.663). The difference in time between the two tasks (30.5 seconds for raw acquisition vs 16 seconds for linear registration) was also significant (W = 29,582, p < 0.001).

For the raw acquisition exercise, trained raters took vastly different times inspecting the training dataset with median times ranging between 15 seconds to 3 minutes (Table 2). Both expert and trained raters generally spent longer before passing an image (median 47 seconds) than failing it (median 41 seconds) (Mann Whitney test, W = 126,898; p = 0.042) (Fig 4), and this delay was more significant when looking only at trained raters (median 64 seconds vs 42.5 seconds; W=87,330, p = 0.005). On the linear registration exercise, raters spent less than a minute before assigning a rating (Table 3). Contrary to the previous task, there was no significant differences in timing between failing or passing an image on the linear registration task: raters took a median of 24 seconds to either pass or fail an image (W = 98,356; p = 0.85). The same pattern was observed looking at only the trainees’ time data (29 seconds for both ratings; W = 58,110, p = 0.420) (Fig 5).

**Table 2.**
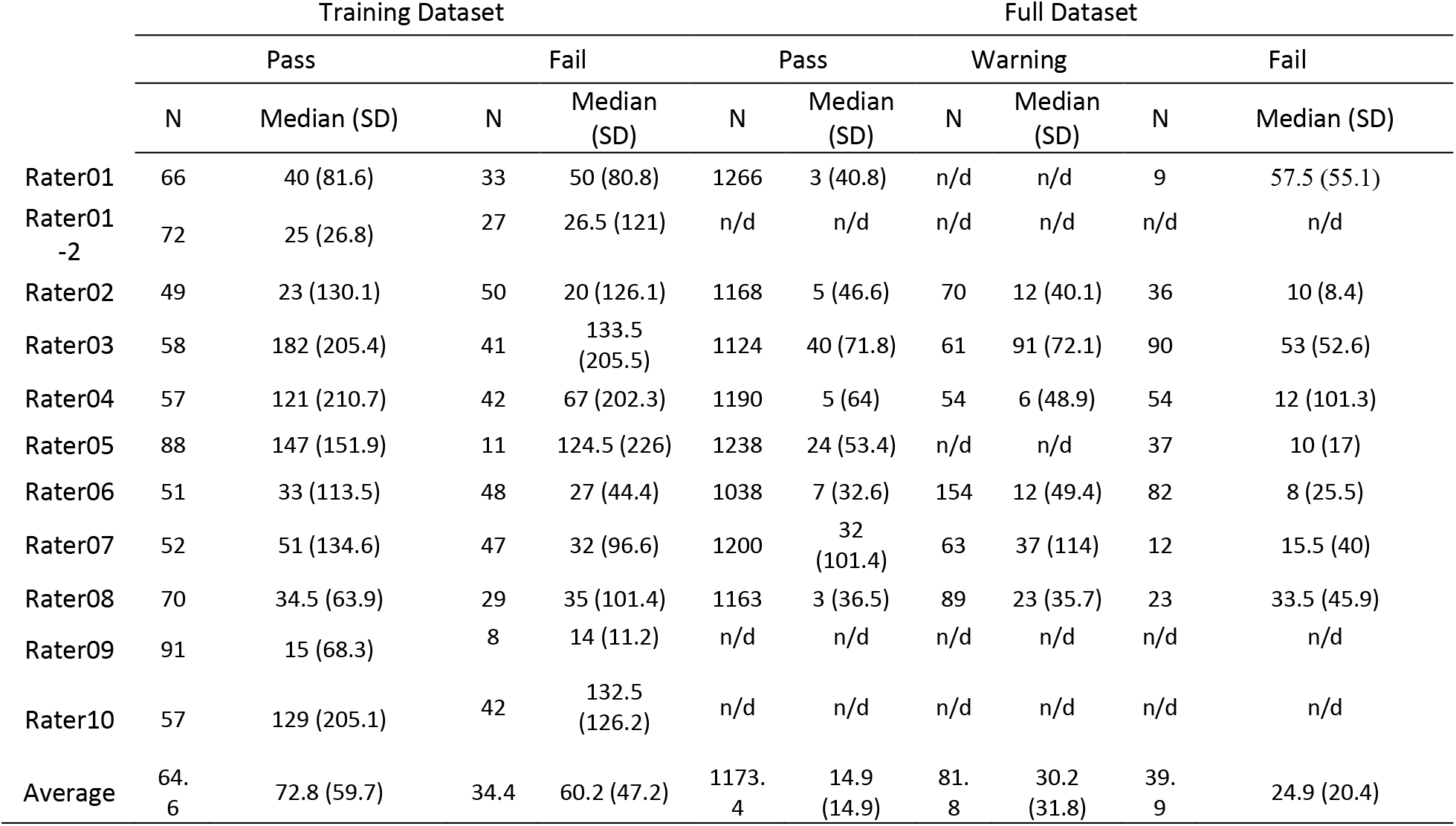
Average and Standard Deviation of the time in seconds spent on an image before assigning a rating by the different raters on the raw acquisition qc exercise. Any time higher than 900 seconds (15 minutes) was assumed to be time spent away from the computer and thus discarded for the timing analyses. (n/d = no data).

|  | Training Dataset |  |  |  | Full Dataset |  |  |  |  |  |
| --- | --- | --- | --- | --- | --- | --- | --- | --- | --- | --- |
|  | Pass |  | Fail |  | Pass |  | Warning |  | Fail |  |
|  | N | Median (SD) | N | Median (SD) | N | Median (SD) | N | Median (SD) | N | Median (SD) |
| Rater01 | 66 | 40 (81.6) | 33 | 50 (80.8) | 1266 | 3 (40.8) | n/d | n/d | 9 | 57.5 (55.1) |
| Rater01-2 | 72 | 25 (26.8) | 27 | 26.5 (121) | n/d | n/d | n/d | n/d | n/d | n/d |
| Rater02 | 49 | 23 (130.1) | 50 | 20 (126.1) | 1168 | 5 (46.6) | 70 | 12 (40.1) | 36 | 10 (8.4) |
| Rater03 | 58 | 182 (205.4) | 41 | 133.5 (205.5) | 1124 | 40 (71.8) | 61 | 91 (72.1) | 90 | 53 (52.6) |
| Rater04 | 57 | 121 (210.7) | 42 | 67 (202.3) | 1190 | 5 (64) | 54 | 6 (48.9) | 54 | 12 (101.3) |
| Rater05 | 88 | 147 (151.9) | 11 | 124.5 (226) | 1238 | 24 (53.4) | n/d | n/d | 37 | 10 (17) |
| Rater06 | 51 | 33 (113.5) | 48 | 27 (44.4) | 1038 | 7 (32.6) | 154 | 12 (49.4) | 82 | 8 (25.5) |
| Rater07 | 52 | 51 (134.6) | 47 | 32 (96.6) | 1200 | 32 (101.4) | 63 | 37 (114) | 12 | 15.5 (40) |
| Rater08 | 70 | 34.5 (63.9) | 29 | 35 (101.4) | 1163 | 3 (36.5) | 89 | 23 (35.7) | 23 | 33.5 (45.9) |
| Rater09 | 91 | 15 (68.3) | 8 | 14 (11.2) | n/d | n/d | n/d | n/d | n/d | n/d |
| Rater10 | 57 | 129 (205.1) | 42 | 132.5 (126.2) | n/d | n/d | n/d | n/d | n/d | n/d |
| Average | 64.6 | 72.8 (59.7) | 34.4 | 60.2 (47.2) | 1173.4 | 14.9 (14.9) | 81.8 | 30.2 (31.8) | 39.9 | 24.9 (20.4) |

**Table 3.**
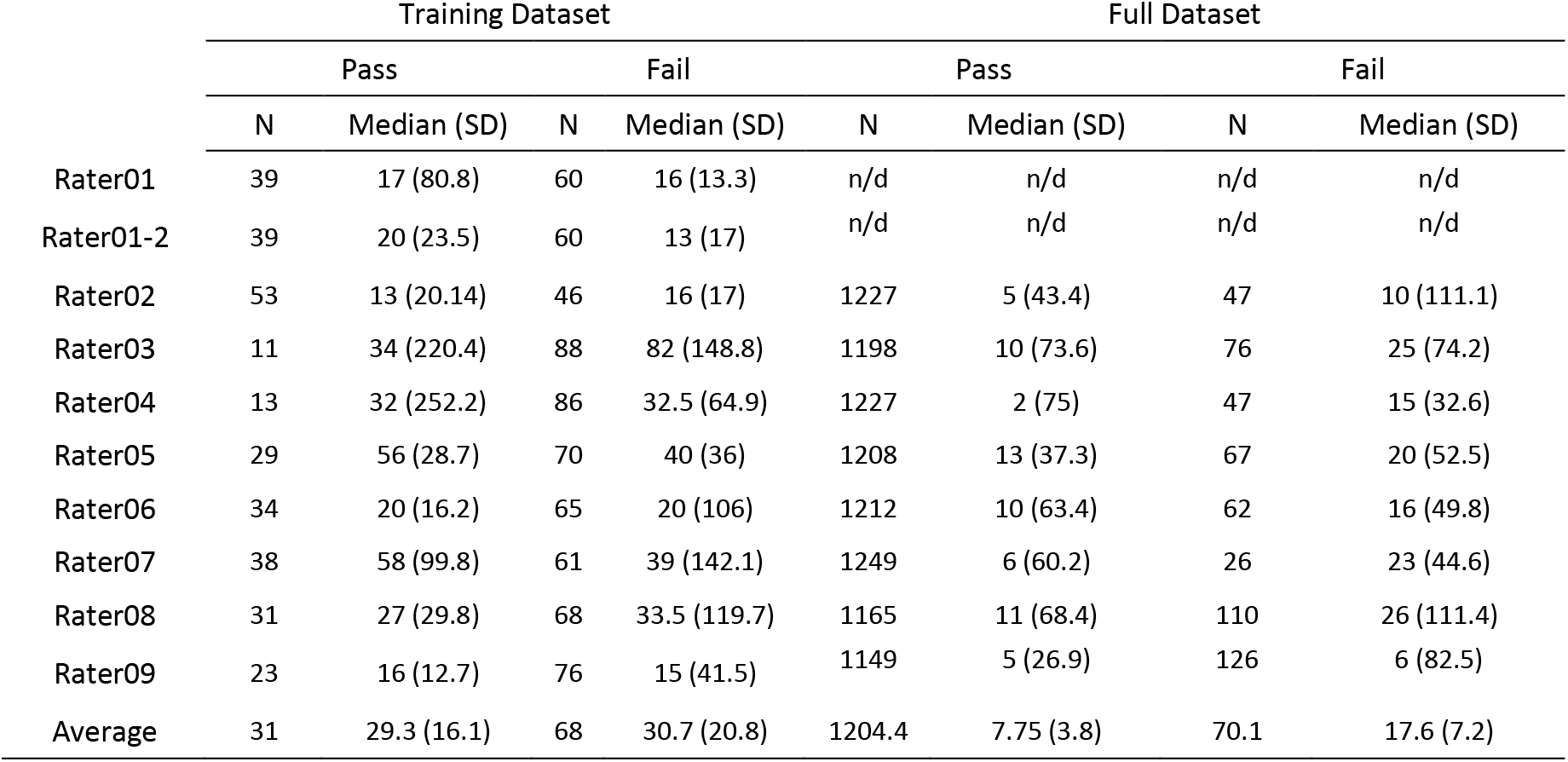
Average and Standard Deviation of the time in seconds spent on an image before assigning a rating by the different raters on the linear registration QC exercise. Any time higher than 900 seconds (15 minutes) was assumed to be time spent away from the computer and thus discarded for the timing analyses. (n/d = no data).

**Figure 4.**
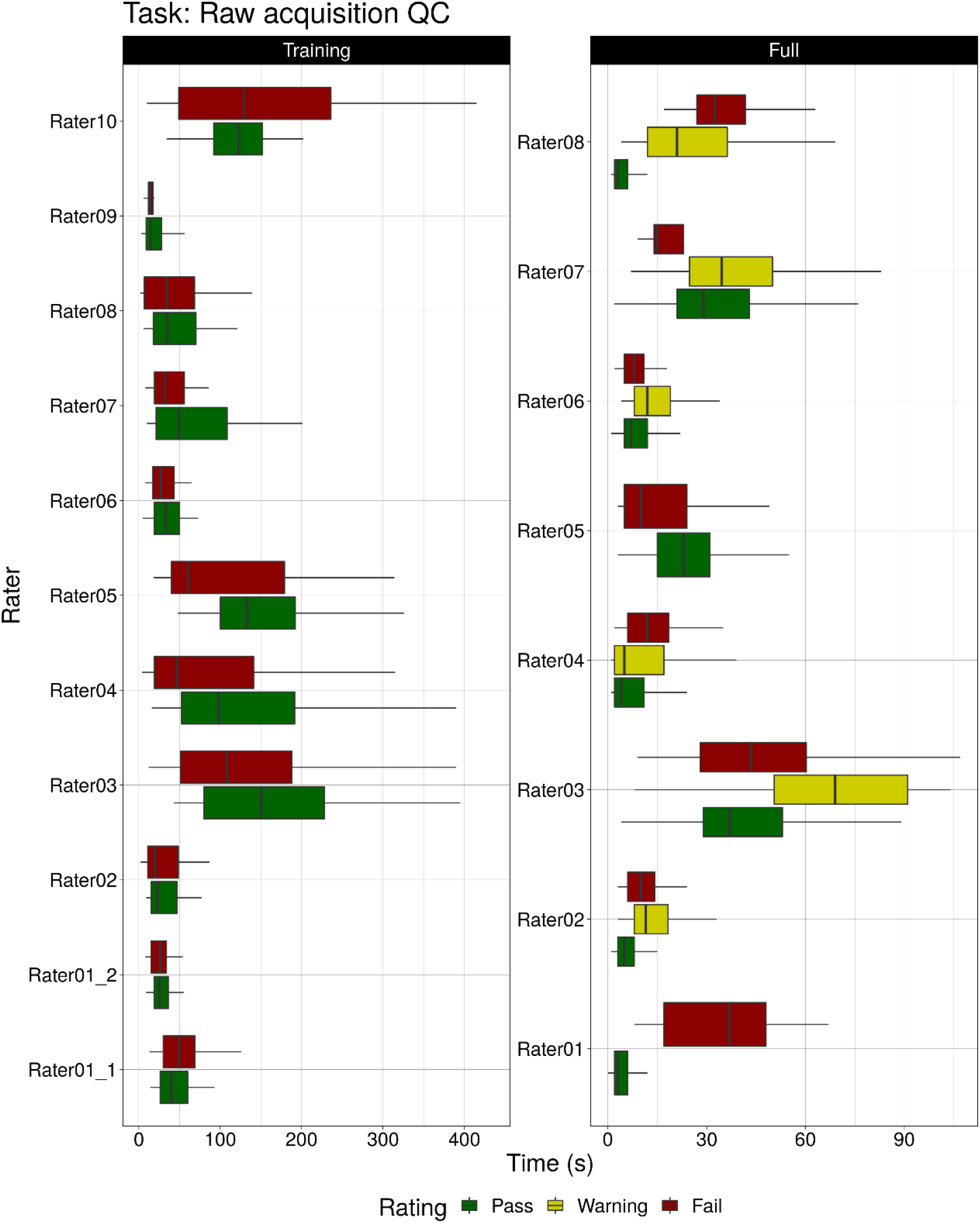
Boxplots represent the time in seconds that the raters spent on the raw acquisition QC images before assigning a rating for the training and full datasets. Raters spend more time passing an image than failing it (Mann Whitney test, W = 126,898; p = 0.042).

**Figure 5.**
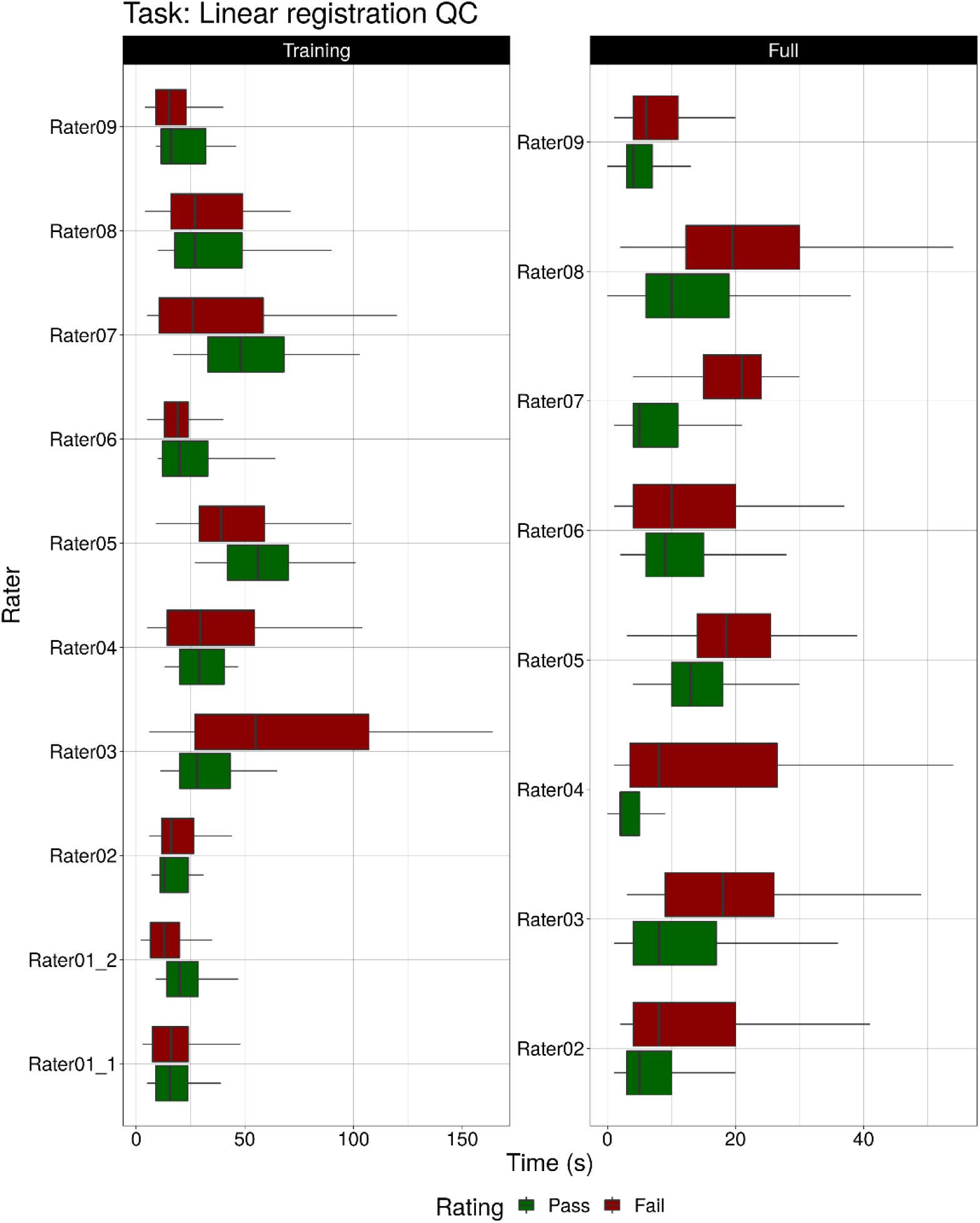
Boxplots represent the time in seconds that the raters spent on the linear registration QC images before assigning a rating for the training and full datasets. No significant difference was found in time taken to PASS or FAIL a dataset.

### Raw acquisition task

A total of 10,196 raw T1w brain images from ADNI3 were divided between 8 raters who went through the QC training protocol. After careful QC inspection, 92% of the of the complete dataset of images was deemed passable according to the QC protocol by the trained raters (PASS = 9,387; WARNING = 491; FAIL = 319), with an acceptance rate ranging from 81 to 99% across the different raters for their respective datasets. Raters took different amount of time depending on the rating given (X2 = 16,743; p < 0.001). The time spent on each image decreased significantly compared to the training dataset (W = 5,204,055; p < 0.001) with the median times between 3 and 91 seconds (Fig. 4, Table 2).

### Registration task

For the final linear registration task, 10,196 QC images were divided between 8 raters who followed the instructions given after the training meetings. They passed 95% of the data (PASS = 9635; FAIL = 561) with acceptance rates ranging from 90 to 98% across raters.

Time varied considerably. Raters took significantly longer to fail an image (16 seconds) than to pass it (8 seconds; W = 1,838,058, p < 0.001). The timing also decreased significantly compared to the training dataset (8 seconds vs 29 seconds; W = 6,193,121, p < 0.001) (Fig. 5). The detailed timing report for all raters can be seen in Table 3.

### Segmentation task

Only 52% (3,674) of the 6,968 output images of the skull segmentation analysis were marked as passing QC by expert Rater 12. This QC task was the fastest, with the expert taking a mean of 3.2 seconds to rate an image. Failing an image was a significantly faster (W = 7,473,831, p < 0.001), taking on average 2.7 seconds compared to the 3.7 seconds for the passing ones.

A second expert rater (Rater 1) reviewed a subset of 1,746 images already rated by Rater 12 with a similar proportion of passing images (54%). The average time for Rater 1 before assigning a rating was 3.9 seconds. Similarly to the other expert, failing an image was significantly faster than passing it (W = 434,559, p < 0.001), taking 3.2 compared to 4.4 seconds on average.

Both expert raters showed an excellent inter-rater agreement (Landis & Koch, 1977) agreeing on 1606 of the 1746 common images yielding a Cohen’s kappa value of 0.83 (Z=35; p < 0.05).

## Discussion

In this article we introduced Qrater, a containerized web-based application with features that facilitate quality control by visual curation of images. Most of these features are accessible via the browser in a user-friendly GUI, which makes Qrater a useful tool for researchers with basic or minimal coding skills. Qrater has proved useful in assisting with QC at the different stages of our analyses process. While several tools intended to aid or automate the QC process have been proposed, given their specificity in QC tasks, MRI modalities or data formats, these tend to be applicable only to a particular set of data and be hard to implement generally. For example, the LONI QC System, a similar web-based system, offers the user to upload their data to their cloud-based service or to install it locally. While using cloud-based computer resources can be convenient sometimes, it also can become unfeasible when dealing with large amounts of data or for datasets with privacy restrictions. The local processing requires the installation of neuroimaging tools. While the generated QC metrics aid significantly (Esteban et al., 2017, 2019), they’re limited to only a fraction of different MRI modalities or QC tasks. Our skull segmentation QC task would be impossible to make use of LONI’s QC System. With Qrater, we wanted to develop a general tool that could adapt to any kind of QC task that depended on the visual inspection of images.

Despite an excellent intra-rater Cohen’s Kappa of 0.81 for the raw QC task and 0.87 for the registration QC task, the expert rater disagreed with himself on 8 datasets for raw QC and 6 datasets for registration QC during the consensus process. The expert even PASSed 3 of the 33 previously selected images deemed as “obvious FAIL”. This demonstrates that despite more than 20 years of QC experience looking at hundreds of thousands of images, expert QC judgement remains subjective and depends on the downstream processing to be completed. Borderline cases will PASS or FAIL depending on the QC rules and on their interpretation by the rater. This not only indicates the need for carefully written visual QC protocols, but strongly justifies the need for objective automatic QC tools.

To evaluate its usability, we tested Qrater on three different example QC tasks from a typical analysis pipeline: data acquisition, image preprocessing and output evaluation. Our goal was to demonstrate the usefulness of Qrater for both experts and raters who went through training for a specific task. Most of the raters who collaborated for the first task of inspecting the MRI acquisition and linear registration tasks went first through the same sessions of training before all evaluating the same pre-selected 99 images datasets. Although some of the Cohen’s kappa values of inter-rater agreement were low between the trained raters, when looking at the agreement with the expert rater on the raw acquisition training task (R01 row in Fig. 3B), only two of the trained raters had a poor agreement (0.24 and 0.3) with the rest being moderate to substantial (0.4 - 0.72) (Portney & Watkins, 2015). These two raters were also among the quickest (Table 2), perhaps indicating that they did not spend enough time doing QC. On the follow-up training meeting, they disclosed that, while they identified artifacts and recorded them in the comments, they PASSed the images thinking that the preprocessing pipeline would be able to deal with them. For the second training exercise of linear registration QC, the inter-rater agreement improved significantly with all but two raters showing substantial to excellent agreement (0.64-0.82) (Portney & Watkins, 2015). The two trainees with poor agreement (0.32 and 0.38) were the most strict, and failed to consider the limitations of linear registration that does not account for residual anatomical variability. These two trainees improved in the follow-up training session before proceeding to the full dataset. These discrepancies underline the need for good, clear training and demonstrates the need for a qualification step to evaluate raters before they begin real QC.

Our inter-rater agreement in this training exercise is comparable to that seen in other similar exercises. Klapwijk, et al. (2019) went through an extensive QC protocol before rating segmentation images by 5 different raters. Only six images (out of 80 scans) were rated in complete agreement and 86% had an agreement by at least three of the five raters. Our QC protocol required our raters to adhere to strict Pass or Fail criteria compared to the 4-level rating scale used by Klapwijk, which might explain their low full agreement compared to ours (29 out of 99 images) on a similar task with a higher number of raters.

Most of the raters in the exercise followed the recommendation of recording qualitative findings as comments independently of the rating given for each image. This aided in the follow-up training session where we went through the images and shared the ratings given and the reasoning. Disagreements were interesting, as the raters who PASSed an image still recorded the same artifacts in the comments. Such qualitative agreement in comments indicates that there is likely a subjective threshold difference between raters rather than a misidentification of the potential artifacts. In the follow-up discussion, several raters mentioned that they passed an image when the artifact found was thought to be mild enough to be corrected by the preprocessing pipeline. Future QC training protocols may need to specifically focus on agreeing on a level of strictness for each QC task to ensure similar thresholds between raters.

Timing varied greatly between the different raters, which was expected given the varying degrees of expertise of the different collaborators. This lack of expertise in QC, the fact that the images were intentionally pre-selected to produce doubt, include a weighted amount of passing and failing images, and the knowledge that their performance would be measured afterwards perhaps explain why this raw acquisition training dataset was slower compared to the other QC tasks.

When the raters went on to review the acquisition and registration of the ADNI3 images, their timing was reduced, as expected. This change was not related to the familiarity with QC in general as it was also present in the expert rater (going from a median 30.5 to 3 seconds per image). This may be because of the expectation that most of the images would pass given that the data has already passed a QC curation within ADNI. With this in mind, there were still some raters who flagged a significant number of images, with the most strict rater flagging close to 20% of their assigned subset. The quality control process should be continuous with evaluation of the quality of the different raters to ensure similarity between them.

Timing and strictness varied between the raters even after the follow-up training session, which is consistent with previous findings of high subjectivity in manual QC protocols despite the training (Esteban et al., 2017; Klapwijk et al., 2019). It’s important to note how halfway through the linear registration exercise with the full ADNI dataset the task was suspended after several raters observed that a high proportion of the images (30-40%) flagged as FAIL when following the instructions given in the training. This contrasted to the 1-5% expected rate of failure of our registration algorithm. The experts met with the trainees in a follow-up meeting and updated the instructions. Specifically, the experts reiterated the importance of considering the limitations of linear registration when dealing with normal anatomical variation. This need to revisit QC instructions demonstrates the importance of having clear PASS or FAIL QC criteria.

Another potential source of variability in QC strictness is the type of post-QC analysis that will be applied to the data. Some neuroimaging analyses are more artifact-forgiving than others, while some methods, like cortical surface extraction, are more affected by artifacts like motion (Bedford et al., 2020). This would potentially modulate a subjective QC threshold.

The third experiment (segmentation) demonstrates the flexibility and efficiency of image-based QC. Generating 3D renderings of the segmented skull surface greatly facilitated the QC task as holes in the surface were easily visible. Such a task would have taken much longer if the rater had to investigate the data on a slice by slice basis. Both expert raters took less than five seconds on most of the images before assigning a rating. Failing an image was faster than passing it. Essentially, as soon as a segmentation error was found, the dataset could be failed without needed to examine all images within the mosaic. Despite the impressive speed of the rating, the inter-rater agreement between both experts was excellent (Portney & Watkins, 2015). Both raters focused on the generated 3D skull surface reconstruction which demonstrates that designing an appropriate QC image can make the specific QC task much easier, faster and more efficient. According to the raters themselves, some of Qrater’s features, like the key-bindings and quick loading times, helped them to breeze through the many images.

The three different experiments, each with a different goal, show the importance of planning beforehand and establishing clear QC criteria to determine if the image rating should be Pass or Fail. In the acquisition experiment, after the follow-up meeting where conflicting images were reviewed and trainees asked QC-related questions that had come up during the first review, the timing decreased considerably. The “Warning” rating in the follow-up raw acquisition exercise, giving the rater the opportunity to flag the dubious images that might warrant re-review made the flagging of the clearly faulty images faster and enables a more consistent QC of borderline cases. The third experiment (segmentation) demonstrates the flexibility and efficiency of image-based QC and the advantage of previous planning. The 3D renderings of the segmented skull surface facilitated the appreciation of holes in the segmented surface. This same task would have taken significantly longer if the raters had to explore the data on a slice-by-slice basis looking through the volume data or by using the common mosaic images. Qrater has vastly improved the way we used to do QC in our laboratory. The instance currently installed on our servers holds more than 100,000 images organized in 48 open and restricted datasets from different research projects with more than 44,000 ratings already recorded without compromising speed or functionality. The latest version of Qrater’s source code is fully available online on GitHub (www.github.com/soffiafdz/qrater).

There are some limitations of Qrater. Qrater functionality depends on the generation of QC images created beforehand outside the application. By leveraging visual inspection of task-specific QC images, Qrater can be applied to many manual QC tasks, but it does not yet take advantage of any automatically extracted quality metrics or automatic prediction of QC status.

Timing information was limited to the time difference between the latest recorded ratings, which could highly bias the data into shorter timings. This indirect measure of time taken to make a QC decision was affected by raters doing a second pass to verify and correct ratings, or detailing comments. All of the rating was done in different personal computers remotely, thus we did not control for internet-speed nor differences in PC hardware quality (monitor definition, size, etc.).

Despite being considered the gold standard, manual QC still has a high cost in time and human resources and remains extremely subjective. Several attempts have been made to both automate and make QC more objective by implementing Machine Learning algorithms to predict the QC label of neuroimaging data are improving, but due to the current performance of these tools they are not a viable replacement for manual efforts for all data (Esteban et al., 2017; Rosen et al., 2018; White et al., 2017). Furthermore, manual QC is still needed for cases where the automatic QC results is not a clear PASS or FAIL.

### Conclusion

In this article we presented Qrater, a novel web-based python application for QC by image curation. From different benchmark tests comprising various tasks done by users with varying amounts of expertise in QC we found that designing and implementing a QC protocol and training that can deal with the subjectivity and high variability among raters is not a trivial task but must be done for reliable QC. We believe that Qrater is a valuable tool that can ease collaboration and facilitate QC, making the task more accessible to beginners and faster for experienced raters.

## Acknowledgments

- This investigation was supported (in part) by (an) award(s) from the International Progressive MS Alliance, award reference number PA-1412-02420
- Data collection and sharing for this project was funded by the Alzheimer’s Disease Neuroimaging Initiative (ADNI) (National Institutes of Health Grant U01 AG024904) and DOD ADNI (Department of Defense award number W81XWH-12-2-0012). ADNI is funded by the National Institute on Aging, the National Institute of Biomedical Imaging and Bioengineering, and through generous contributions from the following: AbbVie, Alzheimer’s Association; Alzheimer’s Drug Discovery Foundation; Araclon Biotech; BioClinica, Inc.; Biogen; Bristol-Myers Squibb Company; CereSpir, Inc.; Cogstate; Eisai Inc.; Elan Pharmaceuticals, Inc.; Eli Lilly and Company; EuroImmun; F. Hoffmann-La Roche Ltd and its affiliated company Genentech, Inc.; Fujirebio; GE Healthcare; IXICO Ltd.; Janssen Alzheimer Immunotherapy Research & Development, LLC.; Johnson & Johnson Pharmaceutical Research & Development LLC.; Lumosity; Lundbeck; Merck & Co., Inc.; Meso Scale Diagnostics, LLC.; NeuroRx Research; Neurotrack Technologies; Novartis Pharmaceuticals Corporation; Pfizer Inc.; Piramal Imaging; Servier; Takeda Pharmaceutical Company; and Transition Therapeutics. The Canadian Institutes of Health Research is providing funds to support ADNI clinical sites in Canada. Private sector contributions are facilitated by the Foundation for the National Institutes of Health (www.fnih.org). The grantee organization is the Northern California Institute for Research and Education, and the study is coordinated by the Alzheimer’s Therapeutic Research Institute at the University of Southern California. ADNI data are disseminated by the Laboratory for Neuro Imaging at the University of Southern California.

## References

Alexander-Bloch, A., Clasen, L., Stockman, M., Ronan, L., Lalonde, F., Giedd, J., & Raznahan, A. (2016). Subtle in-scanner motion biases automated measurement of brain anatomy from in vivo MRI. Human Brain Mapping, 37(7), 2385–2397. https://doi.org/10.1002/hbm.23180

Bedford, S. A., Park, M. T. M., Devenyi, G. A., Tullo, S., Germann, J., Patel, R., Anagnostou, E., Baron-Cohen, S., Bullmore, E. T., Chura, L. R., Craig, M. C., Ecker, C., Floris, D. L., Holt, R. J., Lenroot, R., Lerch, J. P., Lombardo, M. V., Murphy, D. G. M., Raznahan, A., … Chakravarty, M. M. (2020). Large-scale analyses of the relationship between sex, age and intelligence quotient heterogeneity and cortical morphometry in autism spectrum disorder. Molecular Psychiatry, 25(3), 614–628. https://doi.org/10.1038/s41380-019-0420-6

Dadar, M., Fonov, V. S., Collins, D. L., & Alzheimer's Disease Neuroimaging Initiative. (2018). A comparison of publicly available linear MRI stereotaxic registration techniques. NeuroImage, 174, 191–200. https://doi.org/10.1016/j.neuroimage.2018.03.025

Ducharme, S., Albaugh, M. D., Nguyen, T.-V., Hudziak, J. J., Mateos-Pérez, J. M., Labbe, A., Evans, A. C., & Karama, S. (2016). Trajectories of cortical thickness maturation in normal brain development— The importance of quality control procedures. NeuroImage, 125, 267–279. https://doi.org/10.1016/j.neuroimage.2015.10.010

Esteban, O., Birman, D., Schaer, M., Koyejo, O. O., Poldrack, R. A., & Gorgolewski, K. J. (2017). MRIQC: Advancing the automatic prediction of image quality in MRI from unseen sites. PLOS ONE, 12(9), e0184661. https://doi.org/10.1371/journal.pone.0184661

Esteban, O., Blair, R. W., Nielson, D. M., Varada, J. C., Marrett, S., Thomas, A. G., Poldrack, R. A., & Gorgolewski, K. J. (2019). Crowdsourced MRI quality metrics and expert quality annotations for training of humans and machines. Scientific Data, 6(1), Article 1. https://doi.org/10.1038/s41597-019-0035-4

Fonov, V., Evans, A. C., Botteron, K., Almli, C. R., McKinstry, R. C., Collins, D. L., & Brain Development Cooperative Group. (2011). Unbiased average age-appropriate atlases for pediatric studies. NeuroImage, 54(1), 313–327. https://doi.org/10.1016/j.neuroimage.2010.07.033

Fonov, V. S., Dadar, M., Adni, T. P.-A. R. G., & Collins, D. L. (2022). DARQ: Deep learning of quality control for stereotaxic registration of human brain MRI to the T1w MNI-ICBM 152 template. NeuroImage, 257, 119266. https://doi.org/10.1016/j.neuroimage.2022.119266

Gilmore, A. D., Buser, N. J., & Hanson, J. L. (2021). Variations in structural MRI quality significantly impact commonly used measures of brain anatomy. Brain Informatics, 8(1), 7. https://doi.org/10.1186/s40708-021-00128-2

Kim, H., Irimia, A., Hobel, S. M., Pogosyan, M., Tang, H., Petrosyan, P., Blanco, R. E. C., Duffy, B. A., Zhao, L., Crawford, K. L., Liew, S.-L., Clark, K., Law, M., Mukherjee, P., Manley, G. T., Van Horn, J. D., & Toga, A. W. (2019). The LONI QC System: A Semi-Automated, Web-Based and Freely-Available Environment for the Comprehensive Quality Control of Neuroimaging Data. Frontiers in Neuroinformatics, 13. https://www.frontiersin.org/article/10.3389/fninf.2019.00060

Klapwijk, E. T., van de Kamp, F., van der Meulen, M., Peters, S., & Wierenga, L. M. (2019). Qoala-T: A supervised-learning tool for quality control of FreeSurfer segmented MRI data. NeuroImage, 189, 116–129. https://doi.org/10.1016/j.neuroimage.2019.01.014

Landis, J. R., & Koch, G. G. (1977). The measurement of observer agreement for categorical data. Biometrics, 33(1), 159–174.

Monereo-Sánchez, J., de Jong, J. J. A., Drenthen, G. S., Beran, M., Backes, W. H., Stehouwer, C. D. A., Schram, M. T., Linden, D. E. J., & Jansen, J. F. A. (2021). Quality control strategies for brain MRI segmentation and parcellation: Practical approaches and recommendations - insights from the Maastricht study. NeuroImage, 237, 118174. https://doi.org/10.1016/j.neuroimage.2021.118174

Pizarro, R. A., Cheng, X., Barnett, A., Lemaitre, H., Verchinski, B. A., Goldman, A. L., Xiao, E., Luo, Q., Berman, K. F., Callicott, J. H., Weinberger, D. R., & Mattay, V. S. (2016). Automated Quality Assessment of Structural Magnetic Resonance Brain Images Based on a Supervised Machine Learning Algorithm. Frontiers in Neuroinformatics, 10. https://www.frontiersin.org/article/10.3389/fninf.2016.00052

Portney, L. G., & Watkins, M. P. (2015). Foundations of Clinical Research: Applications to Pratice (3rd ed.). F.A. Davis Company.

Power, J. D., Barnes, K. A., Snyder, A. Z., Schlaggar, B. L., & Petersen, S. E. (2012). Spurious but systematic correlations in functional connectivity MRI networks arise from subject motion. NeuroImage, 59(3), 2142–2154. https://doi.org/10.1016/j.neuroimage.2011.10.018

Rosen, A. F. G., Roalf, D. R., Ruparel, K., Blake, J., Seelaus, K., Villa, L. P., Ciric, R., Cook, P. A., Davatzikos, C., Elliott, M. A., Garcia de La Garza, A., Gennatas, E. D., Quarmley, M., Schmitt, J. E., Shinohara, R. T., Tisdall, M. D., Craddock, R. C., Gur, R. E., Gur, R. C., & Satterthwaite, T. D. (2018). Quantitative assessment of structural image quality. NeuroImage, 169, 407–418. https://doi.org/10.1016/j.neuroimage.2017.12.059

Senneville, B. D. de, Manjón, J. V., & Coupé, P. (2020). RegQCNET: Deep quality control for image-to-template brain MRI affine registration. Physics in Medicine &amp$\mathsemicolon$ Biology, 65(22), 225022. https://doi.org/10.1088/1361-6560/abb6be

Weiner, M. W., Veitch, D. P., Aisen, P. S., Beckett, L. A., Cairns, N. J., Green, R. C., Harvey, D., Jack Jr., C. R., Jagust, W., Morris, J. C., Petersen, R. C., Salazar, J., Saykin, A. J., Shaw, L. M., Toga, A. W., Trojanowski, J. Q., & Initiative, A. D. N. (2017). The Alzheimer's Disease Neuroimaging Initiative 3: Continued innovation for clinical trial improvement. Alzheimer's & Dementia, 13(5), 561–571. https://doi.org/10.1016/j.jalz.2016.10.006

White, T., Jansen, P. R., Muetzel, R. L., Sudre, G., El Marroun, H., Tiemeier, H., Qiu, A., Shaw, P., Michael, A. M., & Verhulst, F. C. (2017). Automated quality assessment of structural magnetic resonance images in children: Comparison with visual inspection and surface-based reconstruction. Human Brain Mapping, 39(3), 1218–1231. https://doi.org/10.1002/hbm.23911

Yendiki, A., Koldewyn, K., Kakunoori, S., Kanwisher, N., & Fischl, B. (2014). Spurious group differences due to head motion in a diffusion MRI study. NeuroImage, 88, 79–90. https://doi.org/10.1016/j.neuroimage.2013.11.027

